# Regulation of polyamine interconversion enzymes affects α-Synuclein levels and toxicity in a *Drosophila* model of Parkinson’s disease

**DOI:** 10.1101/2025.03.06.641237

**Authors:** Bedri Ranxhi, Zoya R. Bangash, Zachary M. Chbihi, Zaina Qadri, Nazin N. Islam, Sokol V. Todi, Peter A. LeWitt, Wei-Ling Tsou

**Affiliations:** Department of Pharmacology, Wayne State University School of Medicine; Department of Neurology, Wayne State University School of Medicine; Department of Neurology, Henry Ford Health Systems, Detroit, Michigan

**Keywords:** α-Synuclein, Polyamines, Ornithine decarboxylase 1, Spermine synthase, Spermidine synthase, Spermine oxidase, Spermidine/spermine N1-acetyltransferase 1, Parkinson’s disease

## Abstract

Parkinson’s Disease (PD) is a prevalent neurodegenerative disorder with the accumulation and aggregation of alpha-synuclein (α-Syn) as a central pathological hallmark. Misfolding and aggregation of α-Syn disrupts cellular homeostasis, hinders mitochondrial function, and activates neuroinflammatory responses, ultimately resulting in neuronal death. Recent biomarker research indicated a notable increase in the serum concentrations of three L-ornithine-derived polyamines (PAs): putrescine, spermidine, and spermine, each correlating with the progression of PD and its clinical subtypes. However, the role of PA pathways in PD pathology is poorly understood; it is unclear whether elevated PA concentrations are linked to PD pathology, or whether they represent a secondary effect. In this study, we targeted PAs through RNAi knockdown of different PA-interconversion enzymes (PAIE) in a *Drosophila melanogaster* model of PD that overexpresses human, wild-type α-Syn. Our findings reveal significant impact on both the lifespan and motility of PD-model flies when crucial PAIE, such as ornithine decarboxylase 1 (ODC1), spermidine synthase (SRM), spermidine/spermine N1-acetyltransferase 1 (SAT1), and spermine oxidase (SMOX) are targeted. The overexpression of SAT1 and SMOX in this PD model had positive, enduring effects on fly lifespan. Additionally, we noted significant alterations in ⍺-Syn protein levels when PAIE are either knocked down or overexpressed. These findings underscore the role of PA pathways in PD and their potential targeting to modulate ⍺-Syn levels and mitigate neurodegeneration in PD.

## Introduction

Parkinson’s Disease (PD) is a progressive neurodegenerative disorder affecting millions of mid-life individuals worldwide characterized by decline in motor function (including slowed movements, tremors, and cognitive decline)^1, 2^. Its greatest risk factor is increasing age, although environmental factors and genetics may also play a role^3–5^. The motor disorder of PD involves the degeneration of a specific population of neurons located in the substantia nigra which project to the striatum and generate dopamine^6–8^. While most cases of PD appear sporadic^9^, some cases arise from various gene mutations^5^, the most common being *LRRK2*^10^, *and GBA1*^11^. Additionally, more than 2 dozen gene mutations have been associated with causation or enhanced risk for PD^10, 12^. While PD might have multiple etiologies, a central hallmark of the disease is the pathological accumulation of α-synuclein (α-Syn)^13–15^, a small, soluble protein encoded by the *SNCA* gene. α-Syn plays an essential role in synaptic function^16^ and neurotransmitter release^17^.

In PD, α-Syn undergoes structural changes, misfolding, and aggregation into insoluble fibrils^18^. These α-Syn aggregates – commonly seen in PD brain tissue in circular structures known as Lewy bodies^19^ – accumulate over time in the progressive disease and also in aging individuals as an incidental finding^20^. Abnormal α-Syn aggregates interfere with critical cellular processes, including mitochondrial dynamics^21^, proteostasis^22^, and endo-lysosomal membrane integrity^23^. Ultimately, these processes result in selective neuronal damage and death. Thus, there is a need for mechanistic studies to further investigate disease-related α-Syn aggregation and its role in PD progression.

The aggregation process of α-Syn, driven by elevated α-Syn protein levels, is influenced by both genetic^4, 24–29^ and environmental factors^30–32^. Recent biomarker studies suggest that the concentration of polyamines (PAs) is altered in PD^33, 34^. PAs are critical organic polycations that are highly conserved evolutionarily^35^; they are widely distributed in mammalian cells and are essential for various cellular functions, including cell growth^36^, nucleic acid synthesis^37, 38^, ion transport^37, 38^, and apoptosis^39, 40^. Dysregulation of PA homeostasis can lead to various adverse outcomes^41^, culminating in disease and pathology – multiple reports link altered PA metabolism to various types of cancer^42–44^, cardiovascular disease^45^, and neurodegeneration^46–48^. Serum biomarker studies identified an increase in three L-ornithine (ORN)-derived PAs, putrescine (PUT), spermidine (SPD), and spermine (SPM), in early-stage PD patients, all of which correlated with the progression of PD and its clinical subtypes^34^.

PAs can have a dual role in neurodegenerative diseases, functioning as facilitators that preserve neuronal integrity^49^ and contributors to neuronal damage^50^. The role of PA pathways in PD pathology^51, 52^ is unclear: do elevated PA concentrations worsen PD pathology, or do they represent a secondary effect?

The intracellular homeostasis of ORN, PUT, SPD, and SPM is meticulously maintained through synthesis, degradation, and export^53, 54^. PA biosynthesis converts ORN into PUT and with further incorporation of aminopropyl groups into SPD and SPM through specific PA interconversion enzymes (PAIE)^35^. These include PA anabolic and catabolic enzymes^55^. PA anabolic enzymes, such as ornithine decarboxylase (ODC1), spermidine synthase (SRM), and spermine synthase (SMS), facilitate the biosynthesis of PAs^53, 54^. This process involves a series of decarboxylation reactions followed by aminopropylation^56^. PA catabolism is a more complex process in which PAs are broken down into their precursors and is facilitated by a distinct group of PAIE that include spermine oxidase (SMOX), spermidine/spermine N^1^-acetyltransferase (SAT1), and N^1^-acetylpolyamine oxidase (PAOX)^57^. Catabolic PAIE are involved in acetylation and oxidation processes^58, 59^. Additionally, PAs and their byproducts are shuttled into and out of the cellular environment via Na^+^-independent^60^ and ATP-dependent^61^ PA transporters, such as SLC7A2 and ATP13A3. Investigating PA pathways – including metabolites, interconversion enzymes, and transporters^62, 63^ – may provide a better understanding of PD’s pathogenic mechanisms, as indicated by the serum biomarker^33, 34^ and related findings^51^.

Here, we utilized *Drosophila melanogaster* to investigate the significance of PA pathway perturbation in PD pathology, modelled through the neuronal overexpression of human wild-type α-Syn^64, 65^. Our aim was to determine whether targeted PA metabolism could affect α-Syn stability and impact disease progression. We observed that the regulation of PAIEs significantly affects α-Syn toxicity. We identified the PA catabolic enzymes SAT1 and SMOX as critical factors in PD, as they influenced α-Syn protein levels and its effects in *Drosophila*. Our findings provide novel mechanistic insights into a PD model, using α-Syn pathology as a readout to advance biomarker research and set the stage for PA-targeted therapies.

## Materials & Methods

### *Drosophila* Stocks and Maintenance

Publicly available stocks, including elav-Gal4, GMR-Gal4, UAS-CD8GFP, UAS-α-SynWT, UAS-ODC1_RNAi_, UAS-SRM_RNAi_, UAS-SMS_RNAi_, UAS-SMOX_RNAi_, UAS-SAT1_RNAi_, UAS-PAOX _RNAi_, UAS-ATP13A3 _RNAi_, UAS-SLC7A2_RNAi_, UAS-SLC7A14 _RNAi_, and UAS-Ctrl _RNAi_, were procured from Bloomington *Drosophila* Stock Center (BDSC) or Vienna *Drosophila* Resource Center (VDRC). Gifted stocks used in this study were UAS-Ctrl (y, w; +; attP2) (gift from Jamie Roebuck, Duke University) and sqh-Gal4 (gift from Daniel Kiehart, University of Iowa). Stock numbers and Genotypes of all flies were listed in Table 1. Flies overexpressing UAS-DmSAT1 and UAS-DmSMOX were created in our laboratory. cDNA of DmSAT1 (CG4210) or DmSMOX (CG7737) with an in-line HA epitope tag at the 3’ end were synthesized by Genscript (Piscataway, NJ) and cloned into pWalium10.moe and injected for insertion into site attP2. Genomic DNA was extracted for sequencing to confirm line integrity and identity. Flies were reared in a 25°C incubator, 40% relative humidity with a controlled 12/12hour light/dark cycle diurnal environment. For all Gal4-UAS experiments, controls consisted of the elav-Gal4 line crossed with either y, w; +; attP2 or w^1118^ flies, depending on the specific experiment. To synchronize adult progeny, they were collected within 12 hours post-eclosion over a 48-hour period. Groups of 20 age-and sex-matched flies were promptly placed into narrow polypropylene vials containing 5 mL of standard cornmeal fly medium, supplemented with 2% agar, 10% sucrose, 10% yeast, and appropriate preservatives. Food vials were replaced every two to three days.

**Table 1.**
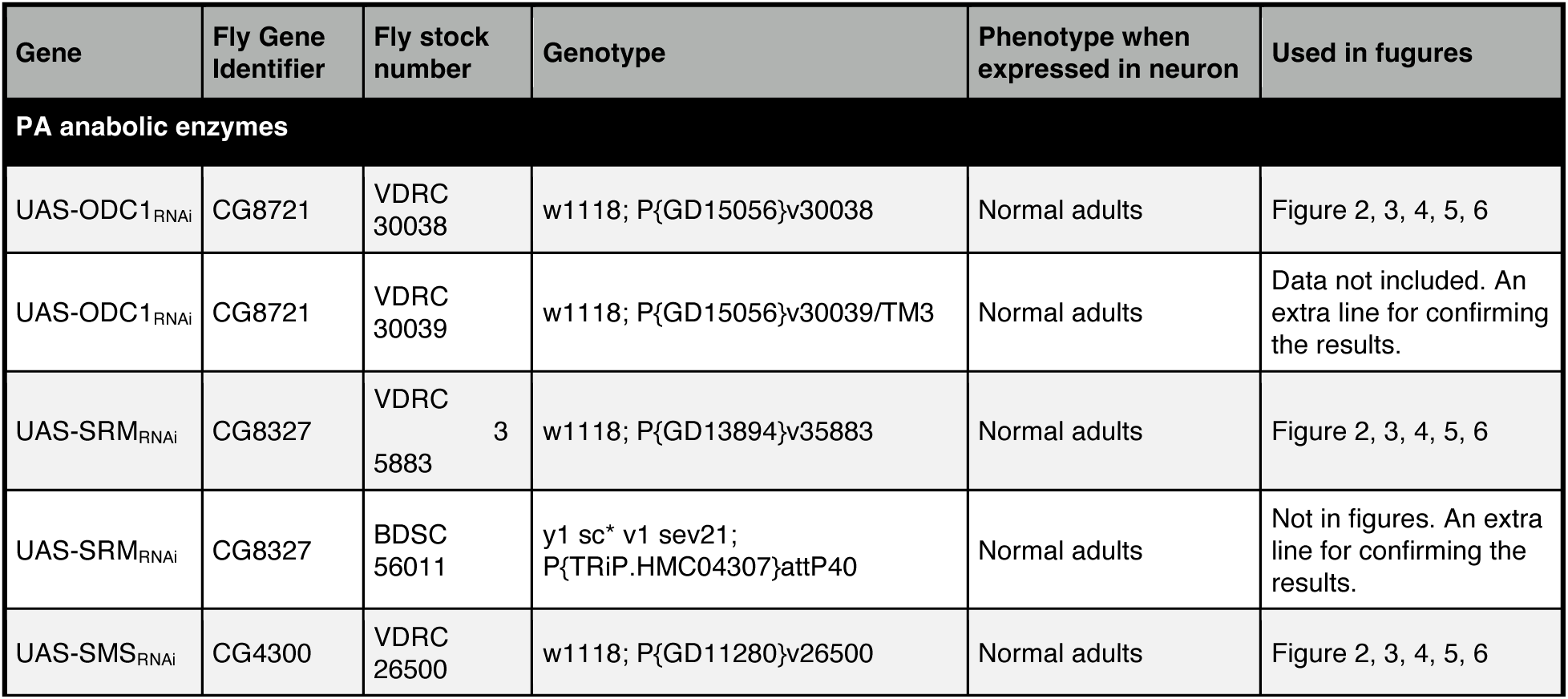
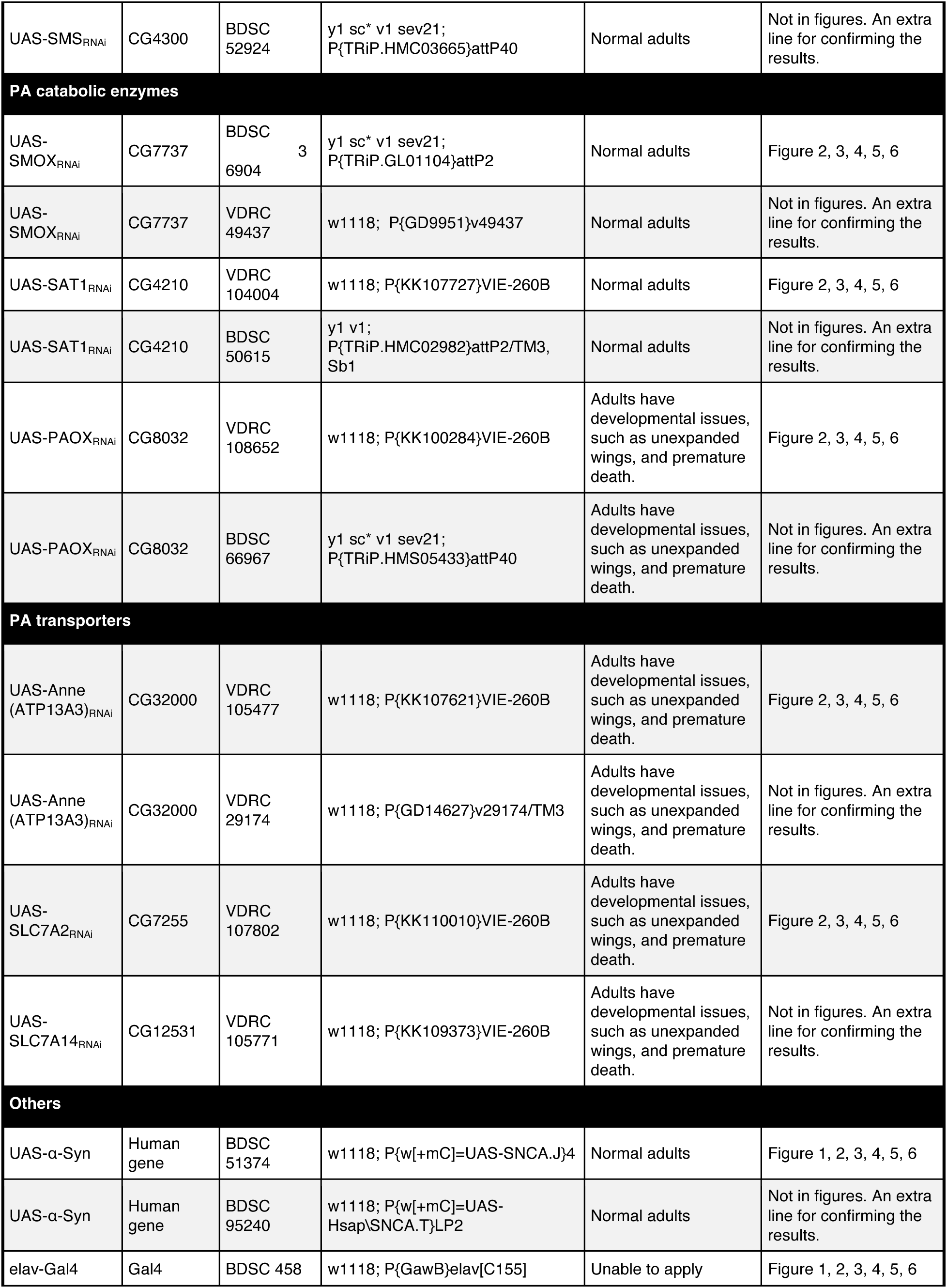
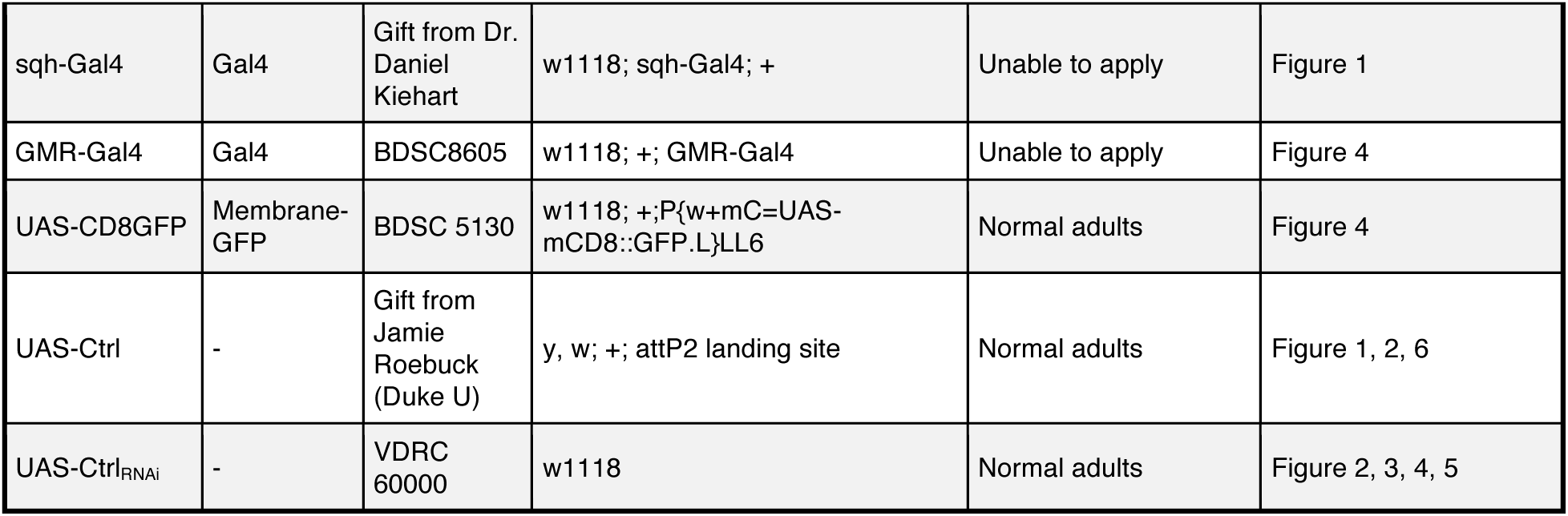
Genotype of flies used in each figure.

### Longevity assay

Approximately 20 adult flies, matched by age and separated by sex within 48 hours of eclosion as adults from their pupal cases, were collected per vial and maintained on standard cornmeal fly medium at 25 °C. Flies were transferred to fresh food vials every 2–3 days, and mortality was monitored daily until all flies had died. Total fly numbers are indicated in each figure. Survival data were analyzed using the log-rank test in GraphPad Prism (San Diego, CA, USA).

### Motility assay

Negative geotaxis was measured through a modified Rapid Iterative Negative Geotaxis (RING) assay^64, 66, 67^ involving groups of at least 100 flies. Vials with 20 flies each were tapped to force them to settle at the bottom, and their climbing responses were captured with photographs taken 3 seconds afterward. Weekly records were maintained of the average performance from five consecutive trials, with flies kept on standard food between tests. Positions of the flies in specified vial zones were quantified as percentages using RStudio (Boston, MA, USA), following the methodology described in our previous study^64^.

### CD8-GFP fluorescence measurements

All flies analyzed in this study were heterozygous for both the driver (GMR-Gal4) and transgenes (UAS-CD8-GFP and UAS-RNAi). Progeny were collected at eclosion and aged for 7, 14, and 28 days. At these intervals, fly heads were dissected and imaged for GFP fluorescence using an Olympus BX53 microscope with a 4X objective and a DP72 digital camera. The fluorescence intensity was quantified using ImageJ, as previously described^68, 69^. Statistical analysis of GFP expression was performed using ANOVA in GraphPad Prism 9 (San Diego, CA, USA). All groups had n≥28 flies.

### Western blots

Fourteen fly heads (7 males, 7 females) per biological replicate were homogenized in hot lysis buffer (50 mM Tris pH 6.8, 2% SDS, 10% glycerol, 100 mM dithiothreitol), sonicated, boiled for 10 minutes, and centrifuged at max speed (13,300 rpm) for 10 minutes at room temperature. Protein lysates from at least three replicates were analyzed by Western blotting using 4–20% Mini-PROTEAN® TGX™ Precast Gels, transferred to 0.2 µm PVDF membranes (Bio-Rad, Hercules, CA, USA), and blocked for 30 minutes in 5% milk/TBST. Membranes were incubated overnight at 4°C with primary antibodies: mouse anti-α-Syn (4B12) (1:1000, Sigma-Aldrich) and anti-HA (1:1000, Cell Signaling Technology), followed by secondary peroxidase-conjugated antibodies (1:5000, Jackson ImmunoResearch) for 1 hour at room temperature. Signal detection used EcoBright Pico/Femto HRP substrates (Innovative Solutions), imaged on a ChemiDoc system (Bio-Rad). PVDF membranes were stained with 0.1% Direct Blue 71 for total protein visualization, and band intensities were quantified using ImageLab software (Bio-Rad).

### Statistics

For Western blots, the levels of α-Syn were normalized to Direct Blue staining and compared against control groups. Prism 9 (GraphPad) was used for data visualization and statistical analyses, with all statistical methods detailed in the figure legends.

## Results

### Human α-Syn overexpression reduces lifespan and impairs locomotor function in Drosophila

To investigate the potential effects of modulating PA pathways in the context of PD, we first tested an α-Syn *Drosophila* model in which human α-Syn is expressed either ubiquitously or specifically in neurons using the Gal4-UAS system. This system facilitates targeted temporal and spatial expression of transgenes. We used sqh-Gal4 and elav-Gal4 drivers to direct wild-type human UAS-α-Syn expression in two distinct patterns: ubiquitously across all tissues or pan-neuronally in all neurons, respectively, spanning development and adulthood. As summarized in figure 1A, ubiquitous expression of human α-Syn resulted in a dose-dependent reduction in overall lifespan in both male and female flies; two copies of the α-Syn transgene caused earlier lethality. A more pronounced dose-dependent effect on lifespan was observed for both males and females when α-Syn expression was specifically targeted to neurons, as illustrated in figure 1B.

**Figure 1:**
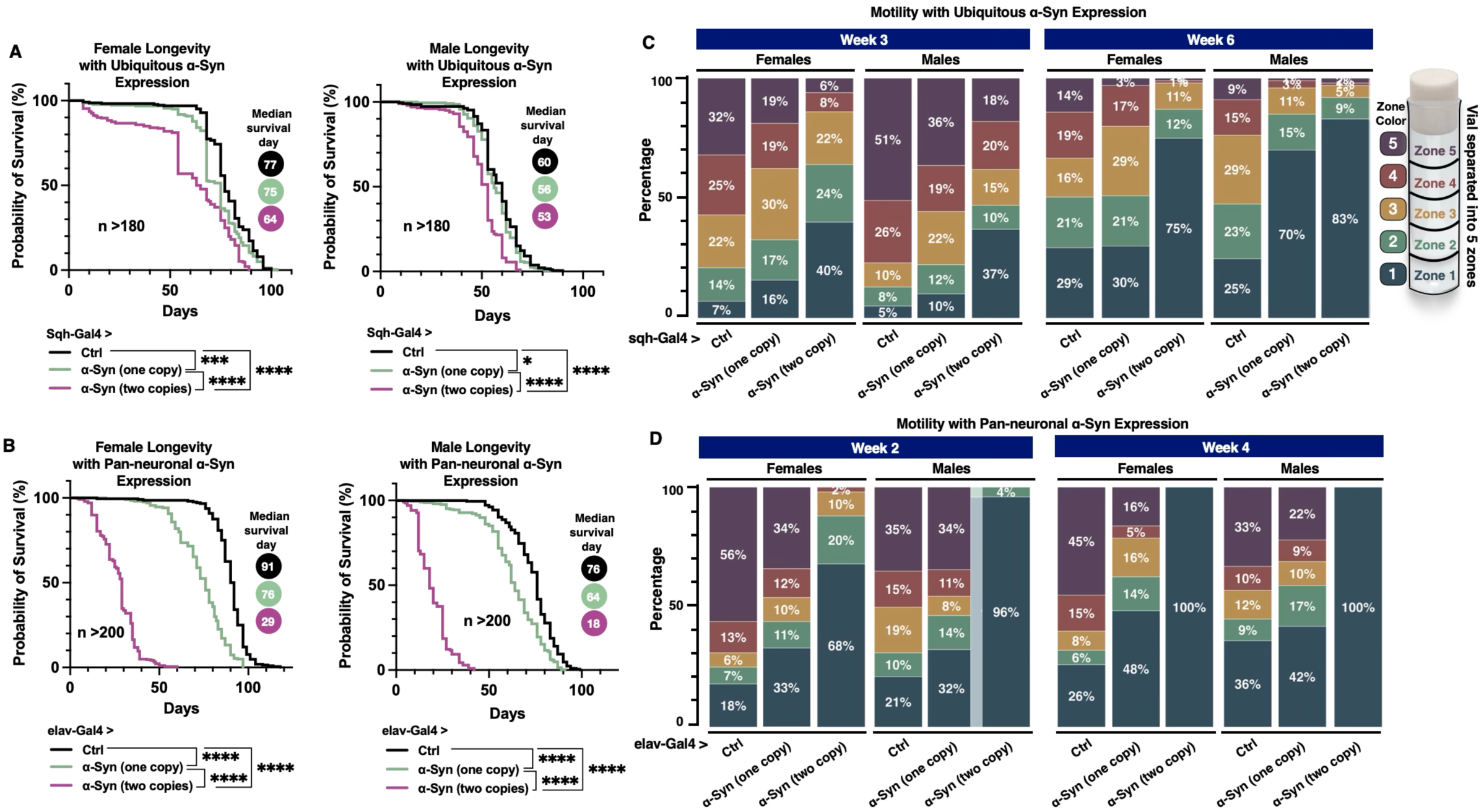
Longevity and motility analyses of flies ubiquitously or pan-neuronally expressing α-Syn. **(A)** Longevity curves of adult female (left) and male (right) flies ubiquitously expressing zero, one, or two copies of α-Syn throughout development and adulthood, using the sqh-Gal4 driver. (**B)** Longevity curves of adult female (left) and male (right) flies expressing α-Syn in all neurons throughout development and adulthood, using the elav-Gal4 driver. Statistical significance for **(A, B)** was determined using log-rank tests: ns (no significance), * (p<0.05), ** (p<0.01), *** (p<0.001), **** (p<0.0001). **(C)** Motility analysis of flies ubiquitously expressing α-Syn, represented as the proportion of flies in each zone of the vial during the RING assay. The left panel shows results at week 3, and the right panel shows results at week 6. Sample size: N ≥ 100 flies per group. **(D)** Motility analysis of flies pan-neuronally expressing α-Syn, represented as the proportion of flies in each zone of the vial during the RING assay. The left panel shows results at week 2, and the right panel shows results at week 4.

Next, we examined a secondary aspect of fly physiology by assessing fly mobility through the Rapid Iterative Negative Geotaxis (RING) assay ^66^. This assay was conducted at three and six weeks post-eclosion of flies ubiquitously expressing α-Syn (figure 1C). At week three, compared to control flies that contained the Gal4 driver in the absence of α-Syn, a smaller proportion of α-Syn-expressing flies reached zones 4 and 5, the highest tiers of the motility index. This decline in motility was also dose-dependent, with flies carrying two copies of the α-Syn transgene exhibiting a more pronounced impairment in both sexes. Notably, sex-specific differences emerged at week six, with a higher proportion of male flies expressing one or two copies of α-Syn remaining in zone 1 compared to their female counterparts, indicating more severe locomotor deficits. These sex-specific differences mirror observations in human populations, where PD is more common in men, with around 65% of patients being male^70, 71^.

We also conducted the same RING assay on flies overexpressing α-Syn pan-neuronally (figure 1D). Due to the early mortality observed in flies with pan-neuronal α-Syn expression, we performed assays at two and four weeks post-eclosion. Pan-neuronal expression of α-Syn led to markedly greater locomotor impairments compared to flies with ubiquitous α-Syn expression. Notably, at both two and four weeks, 96-100% of male flies carrying two copies of the α-Syn transgene remained confined to zone 1 at the bottom of the vial, highlighting the severity of the phenotype. Sex-specific differences were observed as early as week two in flies pan-neuronally expressing two copies of the α-Syn transgene, with the proportion of males remaining in zone 1 being approximately 30% higher compared to females. We conclude that expression of α-Syn in flies leads to reduced motility and longevity in a dose-dependent manner. These baseline data establish the α-Syn overexpression fly model as a valuable tool for investigating the relationship between PA metabolism and α-Syn toxicity.

### Targeting of PAIE modulates α-Syn toxicity in *Drosophila*

Given the elevated levels of PAs reported in the serum of PD patients ^33, 34^, we assessed the effects of PA modulation in human α-Syn-expressing flies. The PA pathway (figure 2A) comprises a complex network of metabolites, including anabolic and catabolic interconversion alongside transport enzymes, that are critical for regulating PA homeostasis. To investigate the effect of PA dysregulation under conditions of α-Syn expression in neurons, we used RNAi-mediated neuronal knockdown of fly orthologs for genes encoding PAIE and PA transporters (figure 2B-Q) and assessed their impact on longevity. We observed that knockdown of ODC1 (figure 2B, 2J), SRM (figure 2C, 2K), or SMOX (figure 2G, 2O), led to a significant increase in the lifespan in both female and male flies. Notably, the knockdown of SAT1 in female flies (figure 2F) but not male (figure 2N) resulted in a reduced lifespan. Neuronal knockdowns of PAOX (figure 2E, 2M), ATP13A3 (figure 2H, 2P), or SLC7A2 (figure 2I, 2Q) caused developmental abnormalities, including unexpanded wings and resulted in markedly earlier lethality in flies with or without α-Syn expression. Due to these developmental issues, we were unable to assess the proper longevity comparison in these conditions. Our results suggest that the lifespan extension observed with ODC1, SRM, or SMOX knockdown highlights the potential protective role of interfering with these PAIE pathways.

**Figure 2:**
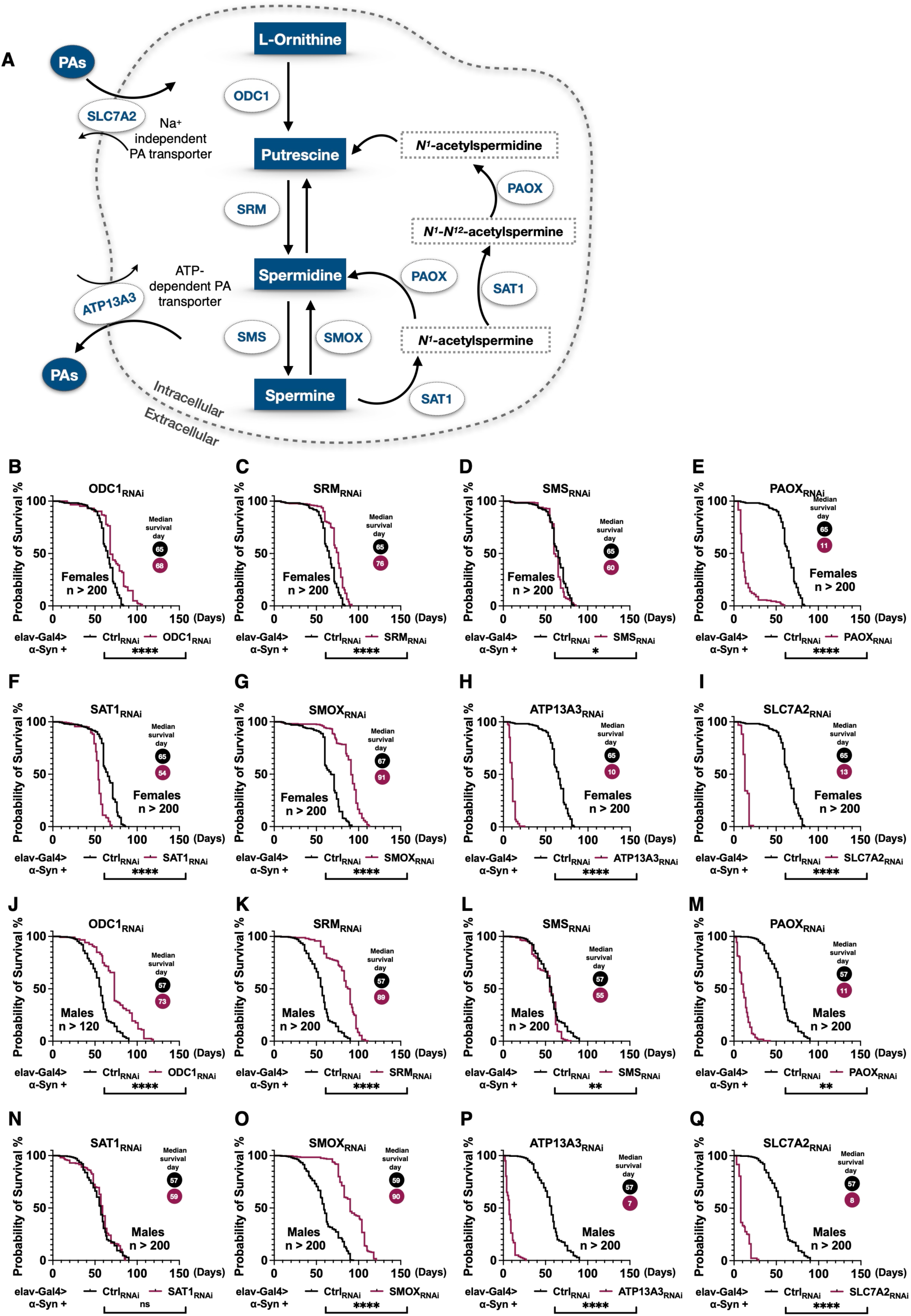
Impact of neuronal knockdown of PAIE on longevity in α-Syn-expressing flies. **(A)** Diagram of the polyamine metabolic pathway, highlighting key enzymes involved in polyamine synthesis, interconversion, and degradation. The pathway includes ODC1, SRM, SMS, SMOX, SAT1, and PAOX, along with the transporters SLC7A2 and ATP13A3. **(B-Q)** Longevity analysis of α-Syn-expressing flies with neuronal knockdown of individual polyamine metabolic enzymes. Panels **(B-I)** depict lifespan data for female flies, while panels **(J-Q)** show lifespan data for male flies. Statistical significance for fly longevity was determined using log-rank tests: ns (no significance), * (p<0.05), ** (p<0.01), *** (p<0.001), **** (p<0.0001).

Next, we examined the impact of PAIE on the motility of α-Syn-expressing flies through the RING assay (figure 3A-F). In week one, we found that knockdown of SMS and SAT1 reduced climbing ability in female flies, with fewer flies reaching the higher zones (4 and 5). Knockdown of SAT1 resulted in 50% of flies remaining in zone 1 (figure 3A). In male flies, suppressing ODC1, SRM, SMOX, or SAT1 initially improved the climbing ability (figure 3B), resulting in a higher percentage of these groups successfully reaching zone 5 and fewer remaining in zone 1. As aforementioned in the longevity results (figure 2), knockdown of PAOX, ATP13A3, or SLC7A2 caused significant developmental issues. These flies exhibited a complete inability to climb and remained in zone 1 (at the bottom of the vial) (figure 3A, 3B).

**Figure 3:**
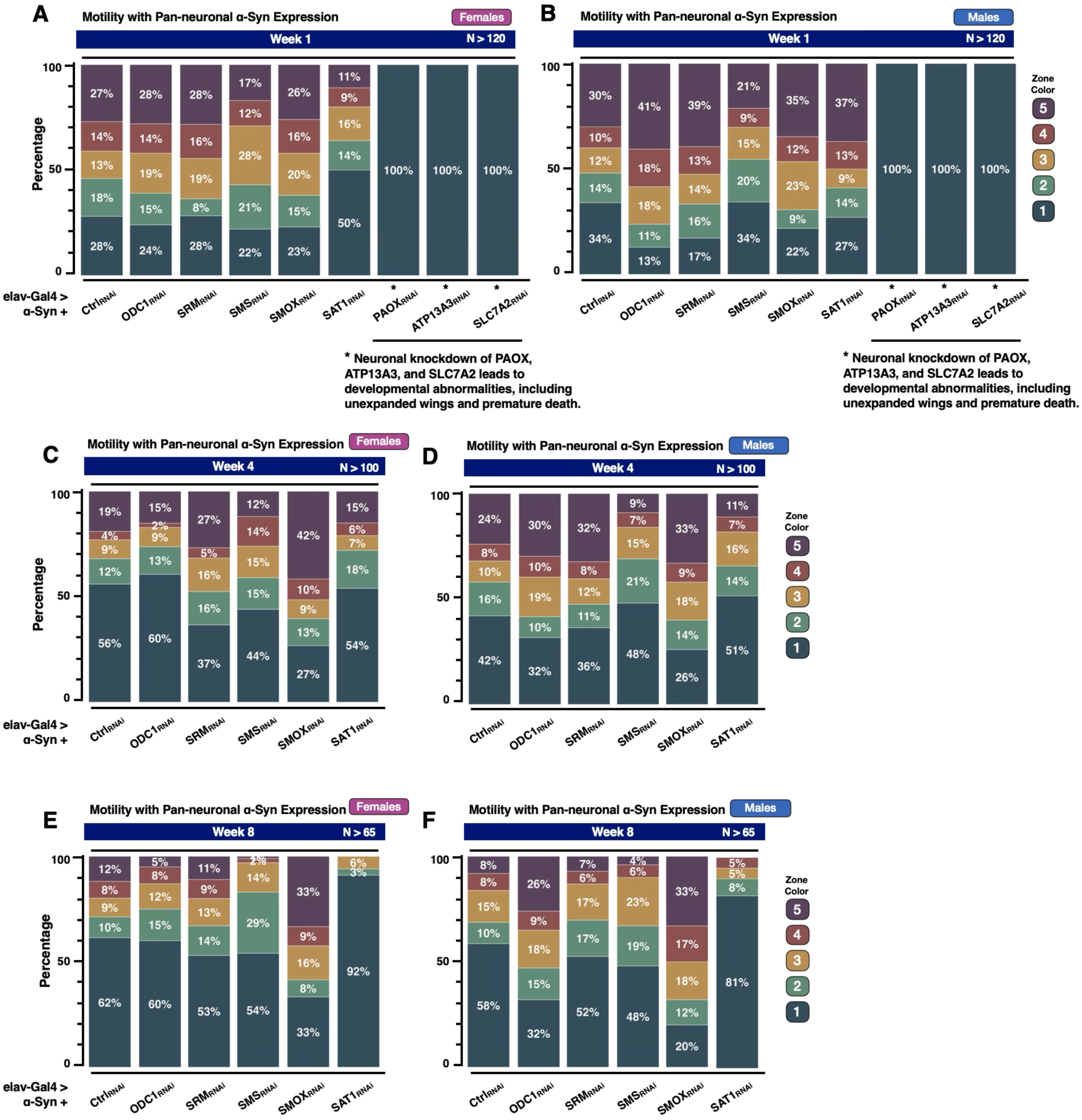
Impact of neuronal knockdown of PAIE on motility in α-Syn-expressing flies. **(A-B)** RING assay results at week 1 for female **(A)** and male **(B)** α-Syn-expressing flies with neuronal knockdown of polyamine metabolic enzymes. **(C-D)** RING assay results at week 4 for female **(C)** and male **(D)** flies. **(E-F)** RING assay results at week 8 for female **(E)** and male **(F)** flies. Motility data for flies with neuronal knockdown of PAOX, ATP13A3, and SLC7A2 are shown only for week 1 due to developmental abnormalities, including unexpanded wings and premature death.

By the fourth week, female flies with knockdown of SRM or SMOX exhibited a marked improvement in mobility, exemplified by the increased proportion of flies reaching zone 5 (figure 3C) and fewer flies in zone 1. In contrast, knockdown of ODC1, SMS, or SAT1 resulted in a decline in female fly mobility, with fewer flies reaching zone 5. In male flies, knocking down ODC1, SRM, or SMOX (figure 3D) resulted in better motility outcomes, as shown by a greater percentage of flies reaching zone 5 compared to the control group. Knocking down SMS or SAT1 caused impaired motility, with fewer flies in zone 5 and more remaining in the lower zones.

In the eighth week, knockdown of SMOX continued to lead to improved mobility, with 33% of both female and male flies (figure 3E, 3F) reaching zone 5, in contrast to 12% and 8% in the control groups, respectively. Interestingly, the knockdown of ODC1 showed an enhancement in motor function exclusively in males (figure 3F), with 26% of the flies successfully reaching zone 5. However, SAT1 knockdown led to decreased mobility in both females and males, with most flies remaining in zone 1 (92% and 81%, respectively). Additionally, the knockdown of SMS hindered fly motility, albeit to a lesser degree for both sexes, with only 1% of female and 4% of male flies reaching zone 5. Our results indicate SMOX knockdown significantly improves locomotor outcomes in the α-Syn model, whereas SAT1 knockdown markedly worsens them.

### PAIE regulates fly eye integrity in the context of α-Syn-Induced Toxicity

To further validate the impact of the PAIE knockdowns on α-Syn toxicity, we tested the *Drosophila* eye, which is a well-established system for studying neurodegeneration and cellular toxicity^68, 72, 73^. We employed a robust assay with the membrane-tagged fluorescent marker CD8-GFP to assess the structural integrity of the fly eyes (figure 4A). In this assay, toxicity manifests as degeneration of internal components within the fly eye ommatidia, resulting in photoreceptor cell loss and a consequent reduction in GFP fluorescence^68^. Enhanced GFP fluorescence indicates improved eye integrity, whereas reduced fluorescence signifies structural degeneration^74^. In our model, the binary Gal4-UAS system drives independent activation of UAS-CD8-GFP, UAS-α-Syn and the PAIE_RNAi_ under the control of the eye-specific GMR-Gal4 driver. In figure 4A, we observed enhanced GFP signal in RNAi-mediated neuronal knockdowns of ODC1, SRM, or SMOX at days 1, 14 and 28. Quantification of GFP intensity (figure 4B-I) confirmed that knockdowns of ODC1 (figure 4B), SRM (figure 4C), or SMOX (figure 4E) at days 14 and 28 led to a notable increase in fluorescence compared to background controls co-expressing CD8-GFP and α-Syn without PAIE gene perturbation. In contrast, flies co-expressing α-Syn with either SAT1_RNAi_ (figure 4F) or PAOX_RNAi_ (figure 4G) at days 14, 28, as well as those with SMS_RNAi_ (figure 4D) at day 28 exhibited significantly reduced GFP intensity compared to non-targeted controls. Moreover, while knockdown of PA transport enzyme ATP13A3 did not alter GFP fluorescence (figure 4H), knockdown of the sodium-independent transporter SLC7A2 led to a significant GFP reduction (figure 4I). Collectively, these findings underscore the significance of PAIE modulation in mitigating α-Syn-induced toxicity, with several key genes, including SMOX, SAT1, ODC1, SRM, and others, play critical roles in regulating neurodegenerative outcomes.

**Figure 4:**
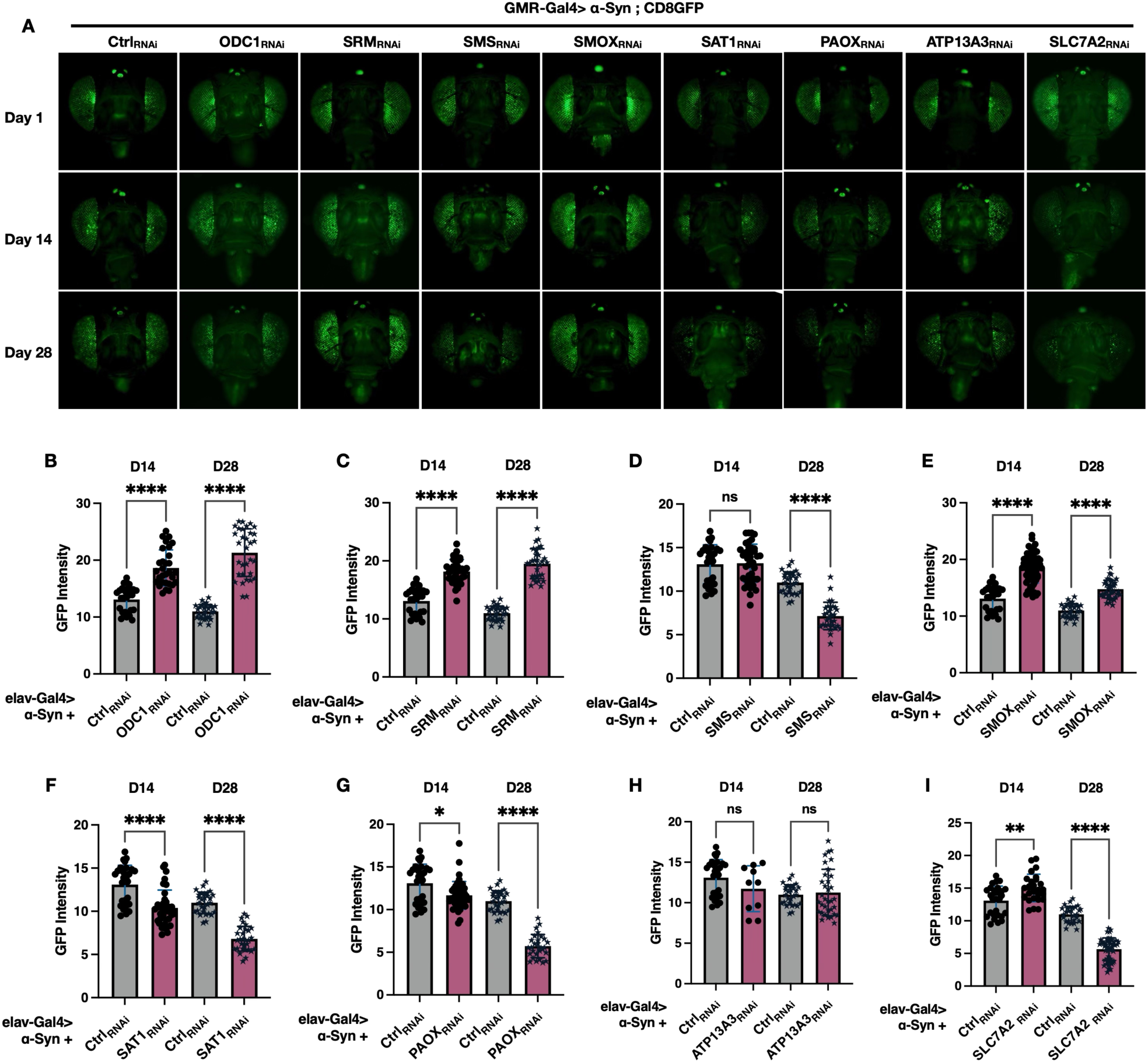
Impact of PAIE knockdown on Drosophila eye integrity in α-Synuclein degeneration. **(A)** CD8-GFP marker visualization of internal eye morphology in female flies at day 1, day 14, and day 28 post-eclosion. **(B-I)** Quantification of GFP fluorescence intensity for internal eye morphology at day 14 and day 28. Sample size: N ≥ 15 per condition. Statistical analysis performed using Brown-Forsythe and Welch ANOVA tests. Significance levels: ns (not significant), * (p<0.05), ** (p<0.01), *** (p<0.001), **** (p<0.0001).

### PAIE knockdowns affect α-Syn protein levels

Encouraged by the marked enhancer and suppressor effects that we observed with the targeting of several PAIEs in various fly tissues and assays, we next evaluated the α-Syn protein levels in flies expressing UAS-α-Syn across all neurons driven by elav-Gal4 (Figure 5A-H, with quantification on the right). Knockdown of the rate-limiting PAIE, ODC1(figure 5A), SRM (figure 5B), as well as the PA transporter, SLC7A2 (figure 5H), did not result in statistically significant alterations in α-Syn protein levels. However, selectively knocking down SMS (figure 5C), ATP13A3 (figure 5D), or SAT1 (figure 5F) resulted in elevated α-Syn protein levels. In contrast, knockdown of PAOX (figure 5E) and SMOX (figure 5G) resulted in attenuated levels of α-Syn. Together, these findings suggest that specific PAIE enzymes differentially regulate α-Syn protein homeostasis, with PAOX and SMOX knockdown reducing α-Syn levels, while knockdown of SMS, ATP13A3, and SAT1 leads to increased α-Syn accumulation.

**Figure 5:**
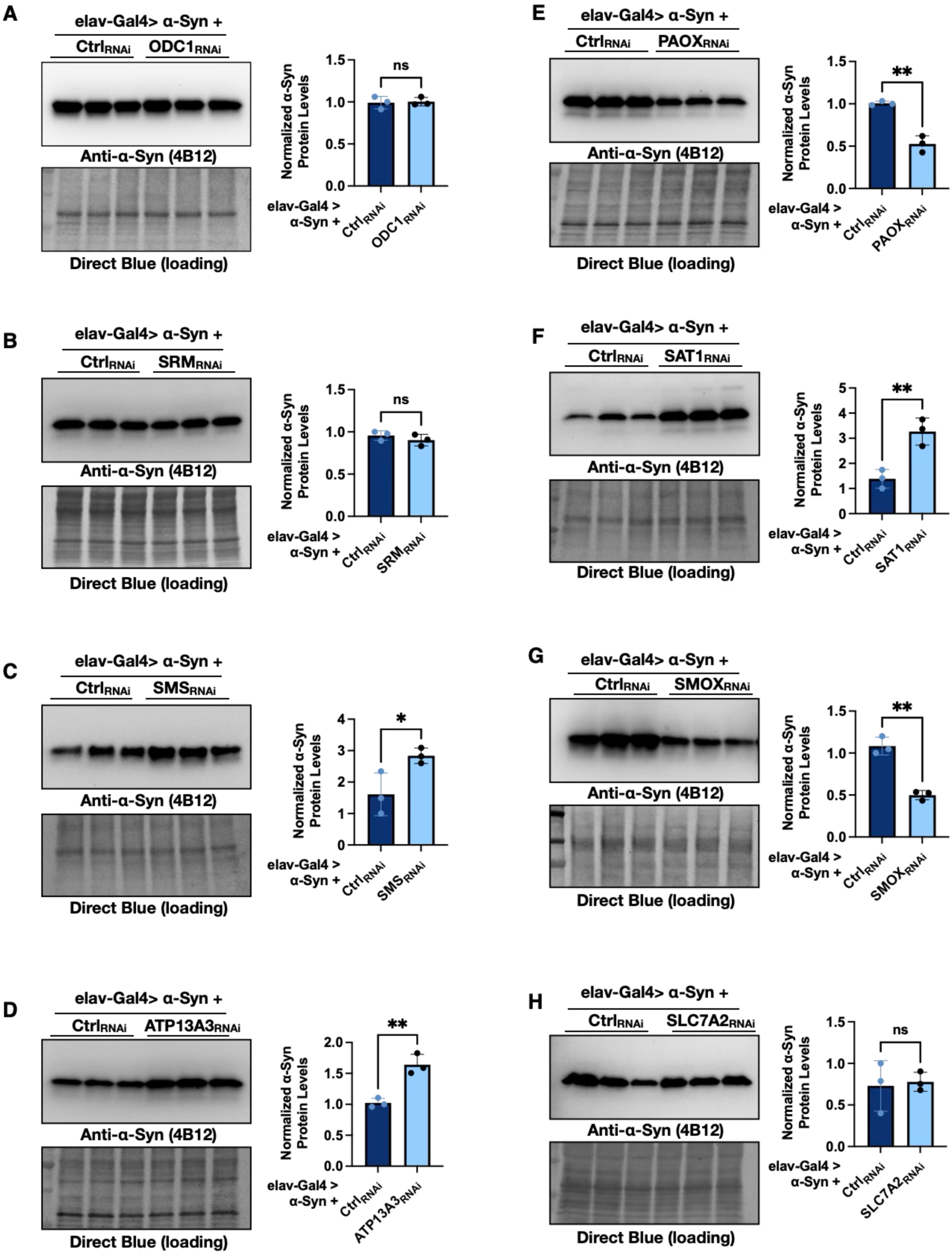
Impact of PAIE knockdown on α-Synuclein protein levels. **(A-H)** Western blot analysis of α-Synuclein protein levels in flies with pan-neuronal expression of α-Syn using the elav-Gal4 driver under conditions of RNAi-mediated knockdown of specific polyamine metabolic enzymes. Statistical analysis was performed using unpaired two-tailed Student’s t-test, Significance levels: ns (not significant), * (p<0.05), ** (p<0.01), *** (p<0.001), **** (p<0.0001).

### Effects of overexpression of SAT1 and SMOX on α-Syn-induced toxicity in *Drosophila*

Intrigued by the effects of SAT1_RNAi_ and SMOX _RNAi_ on longevity, motility, eye integrity, and α-Syn protein levels in the α-Syn *Drosophila* models, we next proceeded to test the effect of individually overexpressing each gene in the presence of α-Syn. We generated new fly lines that carry UAS-DmSAT1 or UAS-DmSMOX by inserting the *Drosophila* SAT1 or SMOX cDNA into the attP2 site on chromosome 3 (figure 6A and 7A, respectively). Pan-neuronal expressing DmSAT1 and α-Syn exhibited a significantly longer lifespan compared to control flies that solely expressed α-Syn (figure 6B). Motility assays (figure 6C) showed that overexpressing DmSAT1 significantly enhances the climbing ability, with females showing a more pronounced improvement than males, reaching the higher zones 4 and 5. Furthermore, Western blot analyses revealed decreased α-Syn levels in the presence of DmSAT1 overexpression (figure 6D).

**Figure 6:**
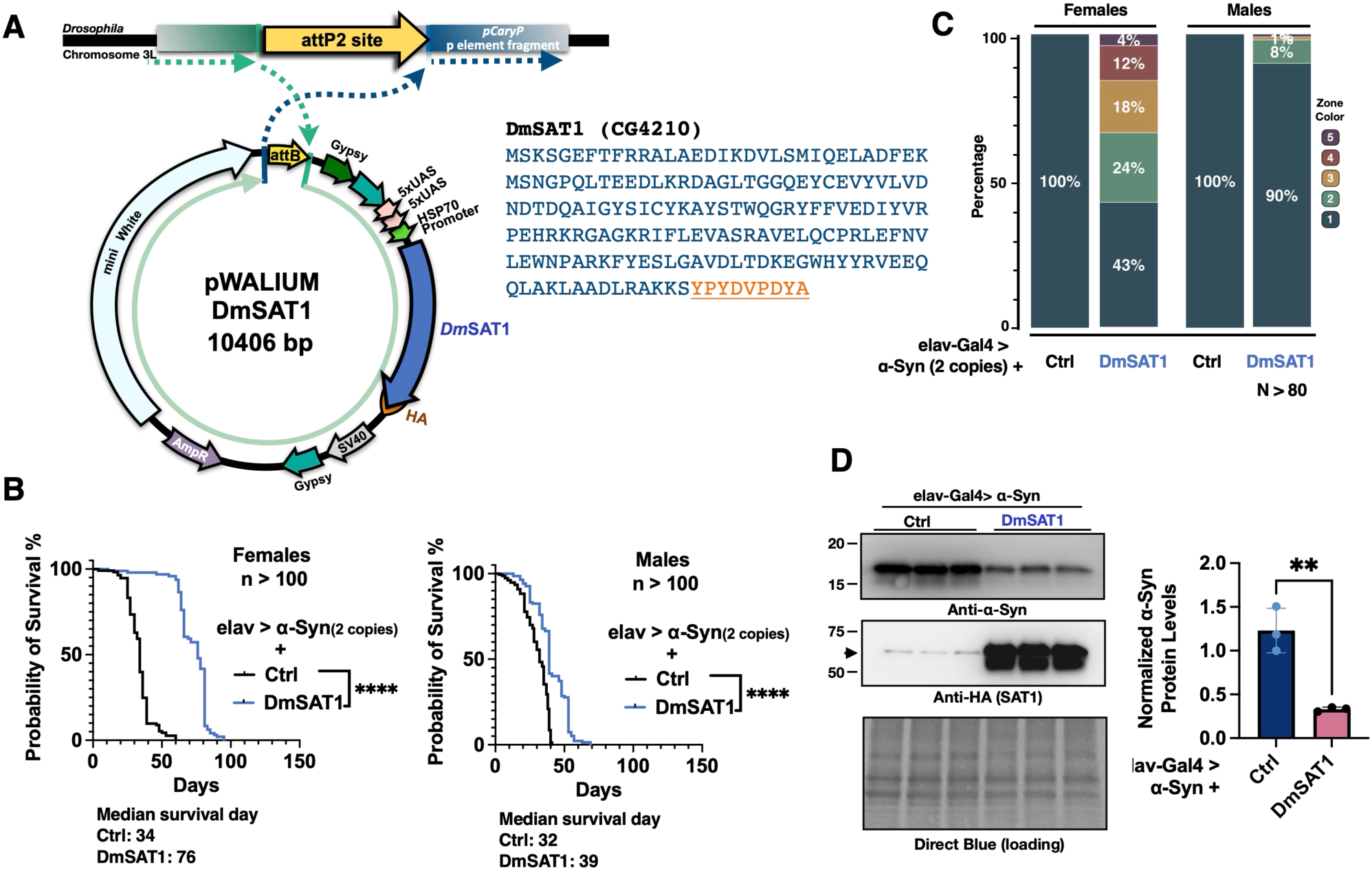
Effects of DmSAT1 overexpression on α-Synucleinopathy. **(A)** Generation of the Drosophila melanogaster (Dm) SAT1 overexpression model. Left: Diagrammatic representation of the cloning strategy used to insert DmSAT1 cDNA into the pWALIUM10.moe vector at the attP2 site. Right: Schematic of the HA-tagged DmSAT1, with the amino acid sequence shown, including the HA tag (underlined). **(B)** Longevity analysis of flies with pan-neuronal expression of two copies of α-Syn, with or without DmSAT1 overexpression. Statistical significance for fly longevity was determined using log-rank tests: ns (no significance), * (p<0.05), ** (p<0.01), *** (p<0.001), **** (p<0.0001). **(C)** Motility analysis using the RING assay of flies with pan-neuronal expression of two copies of α-Syn, with or without DmSAT1 overexpression at week 4. **(D)** Western blot analysis of α-Syn protein levels in flies with pan-neuronal expression of α-Syn, with or without DmSAT1 overexpression. Arrow head: non-specific bands. Statistical analysis for panels. Statistical analysis was performed using unpaired two-tailed Student’s t-test, Significance levels: ns (not significant), * (p<0.05), ** (p<0.01), *** (p<0.001), **** (p<0.0001).

Similarly, we tested flies with pan-neuronal expression of DmSMOX and α-Syn. Intriguingly, DmSMOX overexpression also significantly prolonged the lifespan of α-Syn-expressing flies (figure 7B). RING assays revealed that DmSMOX overexpression improved climbing in both sexes, with males showing a greater increase, as more flies reached zones 4 and 5 compared to females (figure 7C). Western blots also indicated that SMOX overexpression significantly reduced α-Syn protein levels (figure 7D). Collectively, these results suggest that both SAT1 and SMOX overexpression mitigates α-Syn toxicity in *Drosophila* models.

**Figure 7:**
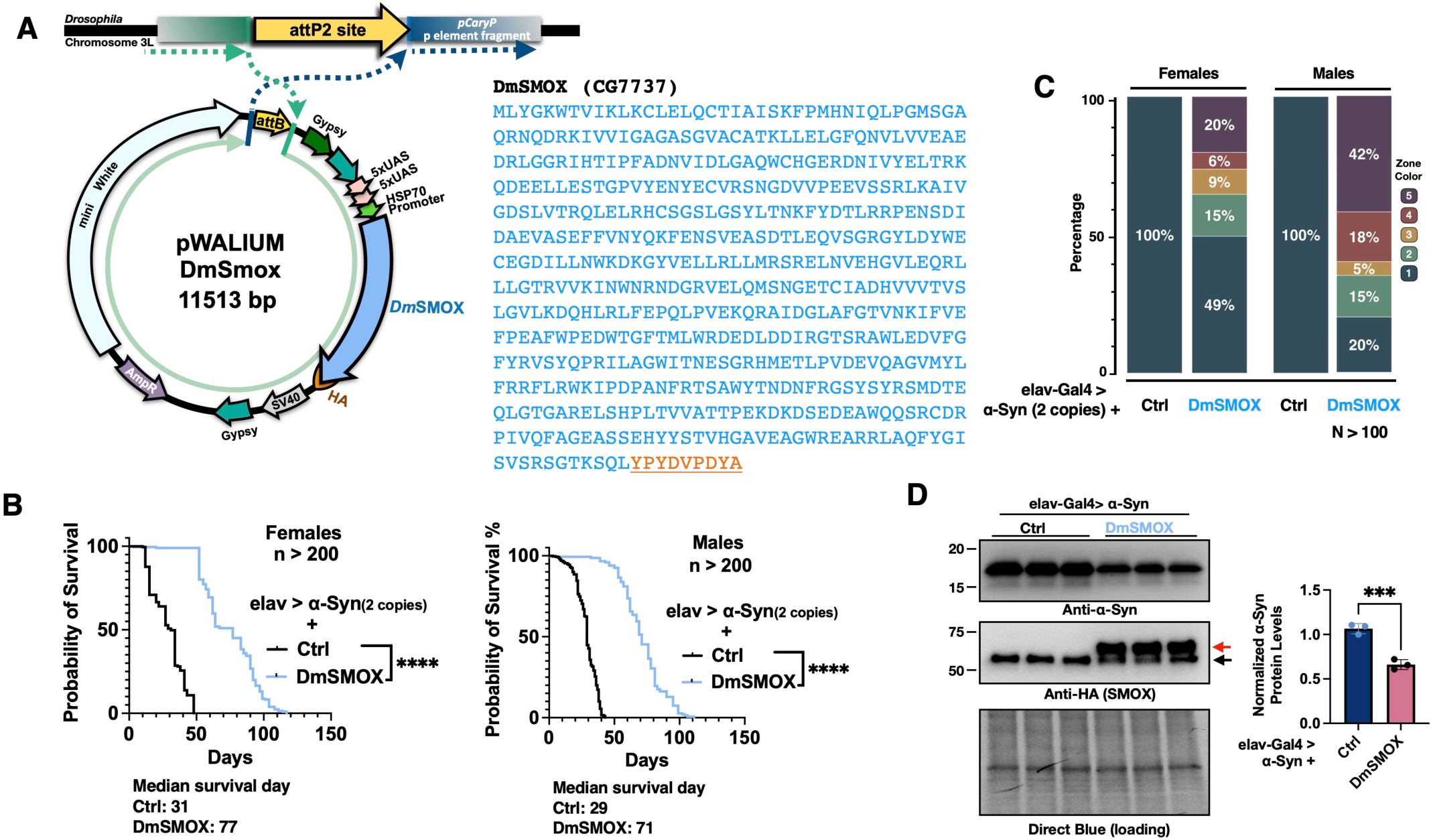
Effects of DmSMOX overexpression on α-Synucleinopathy. **(A)** Generation of the Drosophila melanogaster (Dm) SMOX overexpression model. Left: Diagrammatic representation of the cloning strategy used to insert DmSMOX cDNA into the pWALIUM10.moe vector at the attP2 site. Right: Schematic of the HA-tagged DmSMOX, with the amino acid sequence shown, including the HA tag (underlined). **(B)** Longevity analysis of flies with pan-neuronal expression of two copies of α-Syn, with or without DmSMOX overexpression. Statistical significance for fly longevity was determined using log-rank tests: ns (no significance), * (p<0.05), ** (p<0.01), *** (p<0.001), **** (p<0.0001). **(C)** Motility analysis using the RING assay of flies with pan-neuronal expression of two copies of α-Syn, with or without DmSMOX overexpression at week 4. **(D)** Western blot analysis of α-Syn protein levels in flies with pan-neuronal expression of α-Syn, with or without DmSMOX overexpression. Red arrow: the bands of SMOX. Black arrow: non-specific bands. Statistical analysis for panels. Statistical analysis was performed using unpaired two-tailed Student’s t-test, Significance levels: ns (not significant), * (p<0.05), ** (p<0.01), *** (p<0.001), **** (p<0.0001).

## Discussion

The finding of elevated concentrations of L-ORN-derived PAs in the serum of PD patients ^34^ suggested a potential systemic disruption in PA metabolism. Given the highly-regulated nature of PA homeostasis^75^, we hypothesized that altered concentrations of the various PAs influence PD pathology, as suggested in prior studies^51, 76^. Intrigued by these results, we investigated the functional significance of the PA pathway in an α-Syn-dependent *Drosophila* model of toxicity, focusing on the roles of PAIE and PA transporters. Our findings reveal several unique phenotypic outcomes tied to specific genetic modifications, emphasizing the role of PA pathways in synucleinopathies and their potential as therapeutic targets (Figure 8).

**Figure 8.**
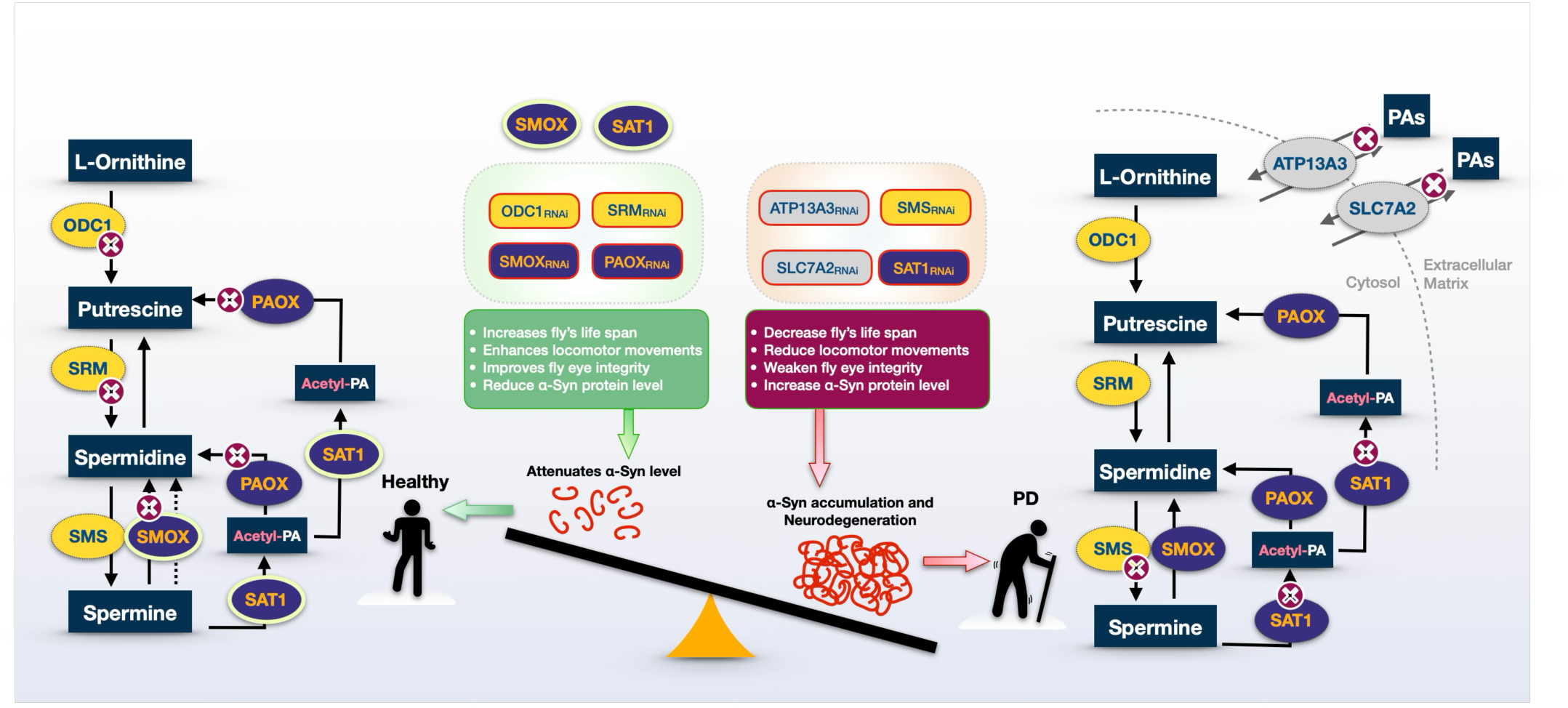
Proposed model of the polyamine pathway and its regulation of α-Synuclein levels and toxicity. Illustration of the PA pathway, emphasizing how regulation of PAIE through RNAi knockdown (rectangles) or overexpression (ovals) affects α-Syn protein levels and toxicity. The model highlights the influence of individual enzymes on polyamine metabolism, α-Syn accumulation, fly lifespan, motility, and eye integrity. The left half represents a beneficial polyamine metabolism, where knockdown of ODC1, SRM, SMOX, PAOX or overexpression of SMOX, SAT1 reduces α-Syn levels and toxicity, promoting health and longevity. In contrast, the right half illustrates the PD-like condition, where knockdown of SMS, SAT1, ATP13A3, or SLC7A2 leads to either α-Syn accumulation or increased toxicity, which leads to neurodegeneration.

In knockdown experiments, toxicity readouts from longevity, motility, and GFP eye integrity assays closely correlated with α-Syn protein levels, highlighting the impact of various PAIE on α-Syn toxicity (figure 8). SMOX knockdown mitigated neurodegeneration by reducing α-Syn protein levels, improving eye integrity, extending lifespan, and enhancing motility. Conversely, SAT1 knockdown exacerbated α-Syn toxicity, leading to increased α-Syn levels, structural degeneration, shortened lifespan, and impaired motility. Suppression of ODC1 and SRM improved longevity and eye integrity, while motility was enhanced in ODC1 knockdown males and SRM knockdown flies before week 4; however, α-Syn protein levels remained unchanged. Knockdown of SMS resulted in increased α-Syn levels, yet only worsened motility and eye integrity, with no significant effect on lifespan. These findings suggest that distinct PAIE enzymes differentially regulate α-Syn toxicity, either directly or indirectly; they highlight the complex role of PAIE enzymes in α-Syn-associated neurodegeneration.

Our finding that both RNAi-mediated knockdown and overexpression of SMOX in neurons reduce α-Syn toxicity by lowering α-Syn protein levels while enhancing longevity and motility in *Drosophila* is intriguing and suggests a complex regulatory role for SMOX in PA metabolism and neurodegeneration. The oxidation of SPM to SPD, catalyzed by SMOX, generates reactive oxygen species (ROS), including hydrogen peroxide (H₂O₂), as a metabolic byproduct^57, 77, 78^. Consequently, reducing SMOX activity may safeguard neurons from α-Syn-induced damage and toxicity via diminished generation of H₂O₂, thereby lessening the effect of oxidative stress. This protective effect may stem from reduced α-Syn protein levels, as indicated by Western blot analysis. The overexpression of SMOX also results in longer fly lives and decreased levels of α-Syn protein, mirroring the effects observed with SMOX_RNAi_. SMOX overexpression may promote the breakdown of SPM into SPD. This outcome seems advantageous from a physiological perspective, since SPD fosters autophagy^79–81^ and cellular repair processes^82^. By leading to more production of SPD, SMOX overexpression may enhance the autophagic flux, aiding in the removal of α-Syn aggregates and improving phenotypic outcomes^83^. This dual observation suggests that both reduced and elevated SMOX activity play a role in α-Syn toxicity, likely through different mechanisms: reduced SMOX activity decreasing ROS and oxidative stress, while increased SMOX enhancing autophagy through boosted SPD production. These distinct pathways can independently or synergistically contribute to reduced α-Syn toxicity, underscoring the complex roles of PAs and the delicate balance necessary in PA metabolism to maintain neuronal health and proteostasis (Figure 8).

Additionally, we found that neuronal knockdown of the PA transporters, ATP13A3 and SLC7A2, as well as the PAIE gene, PAOX, caused severe developmental abnormalities and resulted in early mortality in the flies. These findings indicate that these ATP13A3, SLC7A2, and PAOX are critical for developmental processes beyond roles of merely maintaining PA homeostasis. Similarly, SAT1 knockdown correlated with increased α-Syn-related toxicity. As the rate-limiting enzyme in PA catabolism^84^, SAT1 facilitates the conversion of PAs into forms that can be reintegrated into metabolic pathways, further degraded, or exported from the cell. A reduction in SAT1 levels may disrupt PA flux, leading to the accumulation of higher-order PAs, like SPD and SPM, which can become cytotoxic at elevated concentrations and disturb cellular homeostasis. According to Western blots, the depletion of SAT1 correlates with an increase in α-Syn protein levels, suggesting a link between SAT1 activity and α-Syn regulation. Conversely, SAT1 overexpression in flies expressing α-Syn resulted in an extended lifespan and a reduction in α-Syn protein levels, suggesting that PA acetylation confers a protective effect.

Overall, our research indicates that specific PAIE and PA transporter genes significantly affect phenotypes in an α-Syn-dependent *Drosophila* model of PD (Figure 8). Future investigations are needed to clarify the molecular mechanisms responsible for these effects and to assess the translational possibilities of adjusting PAIE activity for PD therapeutics, as well as the possibility of their use for biomarker purposes in this incurable disease.

## Author Contributions

BR: data curation, software, validation, formal analysis, writing and editing.

ZRB: data curation, validation, formal analysis, investigation, methodology.

ZMC: data curation, validation, formal analysis, investigation, methodology.

ZQ: data curation, validation, formal analysis, investigation, methodology.

NNI: data curation, validation, formal analysis, investigation, methodology.

SVT: conceptualization, data curation, funding acquisition, software, formal analysis, validation, visualization, methodology, and writing and editing.

PAL: conceptualization, funding acquisition, validation, investigation, visualization, methodology, and writing and editing.

W-LT: conceptualization, data curation, software, formal analysis, validation, investigation, visualization, methodology, writing and editing.

## Acknowledgments

This study was partially supported by the Sastry Foundation Endowed Chair in Neurology (PAL) and NIH grant R01NS086778 (SVT). We sincerely thank the Sastry Foundation for their generous support through the Sastry Foundation Endowed Parkinson’s Disease Research Fund at Wayne State University School of Medicine.

## Data and Materials Availability Statement

Fly lines and source data are available upon request. The authors affirm that all data necessary for confirming the conclusions of the article are present within the article and figures. To request data from this study, please contact W-L. T at.

## Human Participants

**No** human participants were involved in this study.

